# The histone code of the fungal genus *Aspergillus* uncovered by evolutionary and proteomic analyses

**DOI:** 10.1101/2022.01.28.477868

**Authors:** Xin Zhang, Roberta Noberini, Tiziana Bonaldi, Jérȏme Collemare, Michael F. Seidl

**Affiliations:** Theoretical Biology & Bioinformatics Group, Department of Biology, Utrecht University, Padualaan 8, 3584 CH Utrecht, the Netherlands; Westerdijk Fungal Biodiversity Institute, Uppsalalaan 8, 3584 CT Utrecht, the Netherlands; Department of Experimental Oncology, IEO, European Institute of Oncology IRCCS, Via Adamello 16, 20139 Milan, Italy; Department of Oncology and Haematology-Oncology, University of Milano, Via Santa Sofia 9/1, 20122 Milano, Italy

**Keywords:** Aspergilli, chromatin, DNA methylation, histone modification, H3K27methylation, mass spectrometry

## Abstract

Chemical modifications of DNA and histone proteins impact the organization of chromatin within the nucleus. Changes in these modifications, catalyzed by different chromatin-modifying enzymes, influence chromatin organization, which in turn is thought to impact the spatial and temporal regulation of gene expression. While combinations of different histone modifications, the histone code, have been studied in several model species, we know very little about histone modifications in the fungal genus *Aspergillus*, whose members are generally well-studied due to their importance as models in cell and molecular biology as well as their medical and biotechnological relevance. Here, we used phylogenetic analyses in 94 Aspergilli as well as other fungi to uncover the occurrence and evolutionary trajectories of enzymes and protein complexes with roles in chromatin modifications or regulation. We found that these enzymes and complexes are highly conserved in Aspergilli, pointing towards a complex repertoire of chromatin modifications. Nevertheless, we also observed few recent gene duplications or losses, highlighting *Aspergillus* species to further study the roles of specific chromatin modifications. SET7 (KMT6) and other components of PRC2 (Polycomb Repressive Complex 2), which is responsible for methylation on histone H3 at lysine 27 in many eukaryotes including fungi, are absent in Aspergilli as well as in closely related *Penicillium* species, suggesting that these lost the capacity for this histone modification. We corroborated our computational predictions by performing untargeted mass spectrometry analysis of histone post-translational modifications in *Aspergillus nidulans*. This systematic analysis will pave the way for future research into the complexity of the histone code and its functional implications on genome architecture and gene regulation in fungi.

**Data Summary:** The predicted proteomes used in this study are publicly available at the JGI (Joint Genome Institute) MycoCosm repository (1); the species names and abbreviations are listed in Supplementary Table 1. To evaluate the completeness of the predicted proteomes and to obtain a species phylogeny, 758 fungal BUSCO (Benchmarking Universal Single-Copy Ortholog) genes were used, and their names are listed in Supplementary Table 2. The fasta, trimmed alignment, and maximum-likelihood phylogenetic tree files can be found in Supplementary Data 1 and 2 deposited at Zenodo (10.5281/zenodo.6586562). The mass spectrometry results mentioned in Supplementary Table 6 are deposited in the PRIDE database with the dataset identifier PXD033478.

**Impact Statement:** Uncovering how fungi regulate gene expression and genome organization is an important step to understand how they control biological processes such as growth or biosynthesis of bioactive molecules. Despite the known importance of chromatin modifications in controlling a plethora of biological processes across eukaryotes, they remain particularly poorly understood in filamentous fungi, even in model organisms like Aspergilli. Our systematic evolutionary analysis provides a robust framework for the functional analysis of chromatin modifications in *Aspergillus* species and other fungi. Our results do not only implicate candidate enzymes to play a role in new chromatin modifications, but they also point at species that have experienced duplications or losses of genes encoding enzymes for specific chromatin modifications. We therefore expect that this work will set the stage for future research into the complexity of the histone code and its functional implications on gene regulation and genome organization in fungi.

## Introduction

Inside eukaryotic cells, the membrane-bound nucleus facilitates the storage and replication of the genetic material as well as the regulation of gene expression (2). Within the nucleus, DNA is arranged as a chromatin fiber formed by repeated nucleosome units consisting of histone protein octamers (two copies of each H2A, H2B, H3, and H4 histones) that are wrapped by 145-147 bp of DNA (3,4). The chromatin structure is regulated by chemical modifications such as methylation of DNA on cytosines (5mC) and adenines (6mA), and different post-translational modifications (PTMs) on the histone proteins, including methylation and acetylation (5,6). The combination of these different modifications, the histone code, influences the recruitment of diverse nucleosome-associated proteins (e.g., DNA binding proteins), which can cause dynamic transitions between the condensed and transcriptionally silent heterochromatin and the accessible and transcriptionally active euchromatin (7–9). Chromatin dynamics is therefore key for organisms to adapt gene expression patterns in response to a variety of developmental and environmental clues.

In the past decades, several enzymes involved in orchestrating chromatin dynamics have been identified and functionally characterized in diverse eukaryotes (5,9,10). These enzymes can be broadly classified into three distinct types: (i) writers that place modifications on nucleosomes or linker DNA; (ii) erasers that remove these modifications; and (iii) readers that first recognize specific modifications and subsequently recruit writers, erasers, or other proteins to the correct genomic location (11). Chromatin modifiers that catalyze the same type of chemical modification often harbor functionally and evolutionarily conserved catalytic domains. For instance, DNA methylation (5mC) is catalyzed by different DNA methyltransferases (DNMTs) which contain a DNA methyltransferase domain, while histone mono-, di-, and tri-methylation are nearly exclusively catalyzed by proteins containing a SET (Su (var) 3-9, E(z) and Trithorax) domain (10). Next to these catalytic domains, the enzymes often contain accessory domains, which can contribute to the overall stability of the enzyme or establish interactions with other proteins (12). Furthermore, these multi-domain proteins, together with other accessory protein subunits, can form multi-protein complexes to find the correct target region witin the genome and modify the exact amino acid residue, collectively contributing to the specific functions of these complexes (13,14). For example, the histone methyltransferase complex PRC2 (Polycomb Repressive Complex 2) is responsible for the mono-, di-, and tri-methylation of histone protein H3 on the lysine at position 27 (abbreviated as H3K27me1/2/3; the same abbreviation principle will be applied below for other histone PTMs) (15–18). Evolutionary analyses have suggested that the key enzymes responsible for chromatin modifications have evolved and siginificantly expanded early in eukaryotic evolution (19). An analysis of 94 gene families with roles in chromatin modifications reveals that 87 out of 94 gene families emerged more than one billion years ago, and 48 were already present in the last eukaryotic common ancestor (19).

Among eukaryotes, the fungal kingdom is remarkably large and diverse, comprising an estimated 1,5-5 million species with various morphologies, life cycles, and ecological niches (20). The most well-studied model fungal species in respect of chromatin modifications is the unicellular yeast *Saccharomyces cerevisiae*, in which 75 histone PTMs on 43 distinct amino acid positions have been reported with histone methylation and acetylation being most abundant (11). In addition to ancient chromatin modifiers, also lineage-specific innovations such as duplications and losses were found in fungi. For example, Dim-2 and RID (Repeat-Induced Point Mutation (RIP)-Defective) are considered to be fungal-specific DNMTs, while DNMT3 that is typically found in animals and plants seems to be absent in fungi (21–23). However, *S. cerevisiae* is not representative of the complete fungal kingdom, and detailed studies on PTMs are needed for other, less studied fungal species.

The fungal genus *Aspergillus* comprises approximately 350 species, including pathogens of humans, animals, and crops, food contaminants, as well as important cell factories for industrial and medical applications (24). Chromatin dynamics influences gene expression in several Aspergilli and regulates growth, sexual development, secondary metabolite biosynthesis, and virulence (11,25–27). Thus far, very few well-known chromatin modifications and their catalytic protein complexes have been identified and studied in few *Aspergillus* species (11). For example, gene knockout of the H3K9 methyltransferase *Clr4* homolog in *Aspergillus fumigatus* reduced histone H3K9me1/3 results in reduction of radial growth and conidia production, as well as delayed conidiation (26). Similarly, CclA, a subunit of the COMPASS (COMplex of Proteins ASsociated with SET1) complex, is essential for H3K4me2/3 in *A. fumigatus*, and the deletion strain showed increased secondary metabolite production and decreased growth (28). The SAGA (Spt-Ada-Gcn5 Acetyltransferase) complex, which is responsible for histone acetylation, has been functionally characterized in *Aspergillus nidulans* (29). Notably, this complex in *A. nidulans* lacks the Sgf11 and Sus1 subunits that have been described in *S. cerevisiae*, suggesting that this complexe evolves in distinct trajectories in different fungi (29). Besides these few examples, we so far lack a complete view of the occurrence and evolution of chromatin modifiers in the *Aspergillus* genus. Here, we report the phylogenetic analyses of core enzymes involved in the deposition and removal of histone PTMs, and assess the conservation of all the subunits belonging to chromatin regulator complexes in 94 diverse Aspergilli. We also performed proteomics analyses to corroborate our computational predictions and reveal a broader spectrum of histone modifications.

## Material and Methods

### Acquisition of predicted proteomes for 109 fungal species

The predicted proteomes of 109 fungal species were retrieved from JGI (Joint Genome Institute) MycoCosm (https://mycocosm.jgi.doe.gov/mycocosm/home) (1) on 24^th^ January 2020. These species comprise 94 different Aspergilli species, 13 Ascomycota species from other genera, as well as two Basidiomycota species (*Coprinopsis cinerea* and *Dendrothele bispora*) (Supplementary Table 1).

### Phylogenetic analyses of chromatin modifiers

Phylogenetic trees for six major chromatin modifiers were generated using their conserved catalytic domains. Catalytic domain HMMs (Hidden Markov Model) of DNA methyltransferase domain (PF00145), histone methylation SET domain (PF00856), acetyltransferase domain (PF00583), histone acetyltransferase MOZ/SAS FAMILY domain (PF01853), histone deacetylase domain (PF00850), and histone demethylase Jumonji C (JmjC) domain (PF02373) were downloaded from the Pfam database (https://pfam.xfam.org). Hmmsearch, which is part of the HMMER package (v3.1b2) (30), with the Gathering Cut-Off threshold (-cut_ga) was used to identify occurrences of these domains in the predicted proteomes of all 109 fungal species considered here. To filter out short and incomplete fragments, we applied a cutoff calculated by 50% length coverage of the query sequence (domain) or the hit. Subsequently, the identified domain sequences were aligned using MAFFT v7.271 (31). TrimAl v1.2 (-gt 0.1) was used to remove positions in the alignments with gaps in more than 90% of the sequences (32). Maximum-likelihood phylogenetic trees were constructed using IQ-TREE v1.6.10 (33,34) using ultrafast bootstrap and Shimodaira–Hasegawa approximate likelihood ratio test (SH-aLRT) (35) with 1,000 pseudo-replicates, and standard model selection to automatically determine the best-fit model. We predicted additional domains in the analyzed proteins using Hmmsearch with all Pfam domain profiles that are present in the Pfam database (v28.0) and visualized them along the phylogenetic trees using iTOL (36). Lastly, additional TBLASTN searches were performed on the NCBI website (https://blast.ncbi.nlm.nih.gov/Blast.cgi) and checked manually to confirm dubious absences in the domain trees.

### Species tree construction

To construct a species phylogeny of the here analyzed fungi, we used BUSCO v4.0.1 (Benchmarking Universal Single-Copy Ortholog assessment tool) (37) to identify the occurrence of 758 fungal single-copy genes (Supplementary Table 2) within the 109 predicted proteomes (Supplementary Table 1). The identified homologs of single-copy BUSCO genes were subsequently concatenated into a single super-alignment. Then, the maximum-likelihood species phylogeny was estimated using IQ-TREE v1.6.10 (33,34) by employing partitioned model selection, which allows selecting a substitution model for each BUSCO gene. Branch supports were obtained using ultrafast bootstrap and SH-aLRT (35) with 1,000 pseudo-replicates separately.

### Chromatin modifier subunits homology search and gene tree construction

We obtained previously studied chromatin modifier complexes from literature, and the associated protein sequences were downloaded from NCBI (https://www.ncbi.nlm.nih.gov) (Supplementary Table 4). For each subunit of the chromatin modifier complexes, homologs in the analyzed 109 predicted proteomes were detected based on sequence similarity searches. First, BLASTP (v2.2.31+) (38) searches were performed, and an e-value threshold of 10^−2^ was applied. Multiple BLAST matches (High-Scoring Pairs: HSPs) of the same query were concatenated if they were aligned with different non-overlapping regions. Only the non-redundant region was retained when HSPs of the same query were overlapping. For these two conditions, the highest e-value of the different matches was used to represent the merged match. Only matches with the e-value lower than 10^−5^ and covering more than 50% of query or subject were retained for the following analyses. If we observed less than ten hits for the BLASTP search after filtering in Aspergilli, we used PSI-BLAST (Position-Specific Initiated BLAST v2.2.31+) (38) with five iterations (-num_iterations 5) using the best Aspergilli hit from the BLASTP as the query; if no Aspergilli match was retained, we used the same query as for the BLASTP searches. The same filter criterion as for the BLASTP selection was applied to the PSI-BLAST hits, and the results of these two homology search strategies were subsequently jointed.

To account for apparent protein absences due to erroneous gene annotation, we tested these potential losses by using TBLASTN (BLAST v2.2.31+) (38) to query protein sequences against the genome assemblies. Matches with e-value lower than 10^−5^ together with 4,500 nucleotides upstream and downstream were extracted, and Exonerate v2.2.0 (39) was used to predict the gene structure (options: protein2genome, extensive search, showtargetgff, showcigar, and showquerygff options were set as true, and “>%ti(%tcb - %tce)\n%tcs\n” option was used to output the complete aligned coding sequence). Transeq v6.6.0.0 (40) was used to translate the coding sequence into protein sequences. Together with the protein sequences identified from BLASTP and PSI-BLAST, all protein sequences from each subunit were retrieved to build a phylogenetic tree (MAFFT, TrimAl, and IQ-TREE settings were the same as for the tree reconstruction using the catalytic domains above).

For each sequence included in the phylogenic trees, the domain organization was identified and visualized as described above. When the domain organization of a *Aspergillius* protein differed from its orthologues, the corresponding gene model was manually checked in the MycoCosm genome browser (1) and corrected. The presence and absence patterns of 83 subunits from 15 different chromatin modifier complexes were visualized using the Seaborn library in Python (41,42). Lastly, additional TBLASTN searches for dubious absences in the outgroup species were performed on the NCBI website as absences could be caused by older and more fragmented assemblies in the JGI database compared to NCBI.

### Histone enrichment and mass spectrometry

Spores of 4- or 5-days old sporulating *A. nidulans* (AnWT pabaA1 veA1, kindly provided by Prof. Joseph Strauss) growing on ME (Malt Extract) agar medium were harvested with 2 mL sterile water and cultured in 50 mL ME liquid medium at 28 °C, 200 rpm for 16 hours. The sample was centrifuged at 8,000 g for 10 minutes to discard supernatant, and germinating spores were resuspended in 25 mL 0.8 M NaCl and centrifuged again. Ten mL protoplasting solution (20 mg/mL *Trichoderma* lysing enzyme and 5 mg/mL driselease in 0.8 M NaCl) were used to resuspend spores, which were then incubated at 30 °C, 100 rpm for 2-3 hours. Protoplasts were retrieved by filtering through sterile Miracloth and were concentrated by centrifugation at 3,000 rpm, 4 °C for 5 minutes. Histone proteins were extracted according to an adapted histone protein extraction protocol (43). Briefly, 10^8^ protoplasts were resuspended in 1 mL Nuclei Isolation Buffer (1×PBS, 0.5 mM PMSF, 5 μM Leupeptin, 5 μM Aprotinin, and 5mM Na-butyrate) with 1% Triton X-100, followed by vigorous pipetting through a 200 μL pipette several times. Then, centrifuged at 2,300 g for 15 minutes at 4 °C, discarded the supernatant, and resuspended the nuclear pellet in 200 μL Nuclei Isolation Buffer with 0.1% SDS and 250 U Pierce™ Universal Nuclease and incubated at 37 °C for 30 minutes to digest nucleic acids. The protein concentration was measured using the Bradford Protein Assay (44).

The samples were separated on a 17% SDS-PAGE gel and histone bands were excised, chemically acylated with propionic anhydride and in-gel digested with trypsin, followed by peptide N-terminal derivatization with phenyl isocyanate (PIC) (45). Chemical acylation of lysines (which occurs on unmodified or mono-methylated residues) impairs trypsin cleavage, resulting in proteolityc cleavage at arginine residues only. This treatment allows obtaining histone peptides of proper length for MS analysis. The samples were then desalted on handmade StageTips columns (46). Peptide mixtures were separated by reversed-phase chromatography on an EASY-Spray column (Thermo Fisher Scientic), 25-cm long (inner diameter 75 µm, PepMap C18, 2 µm particles), which was connected online to a Q Exactive Plus instrument (Thermo Fisher Scientific) through an EASY-Spray™ Ion Source (Thermo Fisher Scientific), as described (45).

The acquired RAW data were analyzed using the integrated MaxQuant software v.1.6.10. The Uniprot UP000000560 database was used for identification of *A. nidulans* histone peptides. Enzyme specificity was set to Arg-C. The estimated false discovery rate (FDR) was set at a maximum of 1%. The mass tolerance was set to 6 ppm for precursor and fragment ions. Two missed cleavages were allowed, and the minimum peptide length was set to 4 amino acids. Min. score for modified peptides and min. delta score for modified peptides were set to 1. Variable modifications included lysine propionylation, monomethylation + propionylation, dimethylation, trimethylation and acetylation. N-terminal PIC labeling was set as a fixed modification (45). To reduce the search time and the rate of false positives, with increasing the number of variable modifications included in the database search, the raw data were analyzed through multiple parallel MaxQuant jobs (47), setting different combinations of variable modifications. Peptides identified by MaxQuant with Andromeda score higher than 50 and localization probability score higher than 0.75 were quantitated, either manually or by using a version of the EpiProfile 2.0 software (48) adapted to the analysis of histones from *A. nidulans*. Identifications and retention times were used to guide the manual quantification of each modified peptide using QualBrowser version 2.0.7 (Thermo Fisher Scientific). Site assignment was evaluated from MS2 spectra using QualBrowser and MaxQuant Viewer. Extracted ion chromatograms (XICs) were constructed for each doubly charged precursor, based on its *m/z* value, using a mass tolerance of 10 ppm. For each histone modified peptide, the % relative abundance (%RA) was estimated by dividing the area under the curve (AUC) of each modified peptide for the sum of the areas corresponding to all the observed forms of that peptide (49). The AUC values are reported (Supplementary Table 6) and visualized using GraphPad Prism.

## Results and discussion

DNA methylation, histone methylation, and histone acetylation are among the most studied chromatin modifications in animals, plants, and fungi (50–52). Protein complexes involved in writing, reading, and erasing these modifications have been reported in few model fungal species to date (16,22), but the evolution and conservation of these complexes in diverse fungal species have not been largely systematically explored. Besides some research on DNA methyltransferases (21,23) and SET domain-containing proteins (17) that have been performed throughout the fungal kingdom, we still lack the evolutionary overview of other chromatin modifiers and in depth analyses for important fungal groups. To start uncovering the occurrence and evolutionary histories of chromatin modifiers involved in these major modifications in the fungal genus *Aspergillus,* we first focused on six conserved catalytic domains reported to be involved in histone modifications: DNA methyltransferase domain for DNMTs, SET domain for histone methyltransferases (HMTs), acetyltransferase domain and MOZ/SAS FAMILY domain for histone acetyltransferases (HATs), histone deacetylase domain for histone deacetylases (HDACs), and JmjC domain for histone demethylases (HDMTs). We identified homologs harboring these conserved domains in 109 predicted proteomes; the completeness of the predicted proteomes was generally high as 97 had more than 97% complete BUSCO predicted proteins (Supplementary Table 1). The 94 Aspergilli species cover 18 recognized sections: *Polypaecilum*, *Cremei*, *Restricti*, *Aspergillus*, *Cervini*, *Clavati*, *Fumigati*, *Candidi*, *Circumdati*, *Janorum*, *Terrei*, *Flavi*, *Ochraceorosei*, *Bispori*, *Cavernicolarum*, *Usti*, *Nidulantes*, and *Nigri* (53–59), and 15 other species were used as outgroups (Supplementary Figure 1). Two of the outgroup species are Basidiomycota and the remaining species cover each class in the Ascomycota phylum, which provides a robust framework for our evolutionary analyses within Aspergilli. The outgroup species include well-studied model species, some of which are important for crop production, food fermentation, and antibiotics production (60,61). We used the identified homologs for each catalytic domain to determine their evolutionary history using maximum-likelihood phylogenies (Figure 1–4 and Supplementary Figure 2 and 3).

**Figure 1.**
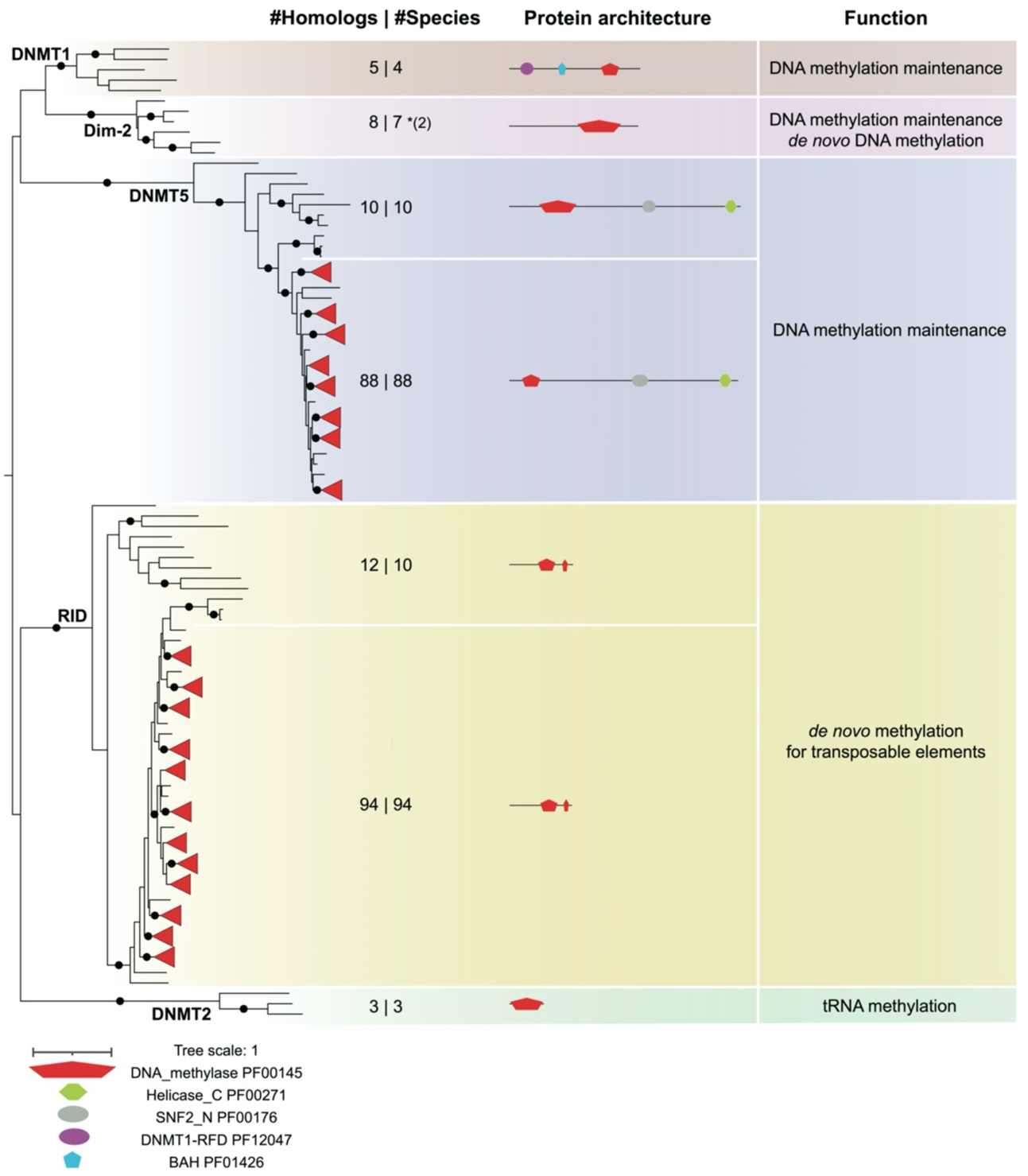
Conservation of DNA methyltransferase families in the *Aspergillus* genus and outgroup species. The maximum-likelihood phylogeny of DNA methyltransferase domain (PF00145) found in 94 Aspergilli and 15 outgroup species was determined using IQ-TREE (37,38). The black dot on the branch indicates ultrafast bootstrap values over 95 and SH-aLRT values over 80. Leaves from Aspergilli with trustworthy support are collapsed and shown as red triangles. The numbers of hits and species are separated by the pipe in the ‘#Homologs | #Species’ column. The number in parentheses after * indicates the additional hits found by the TBLASTN search on NCBI. Additional conserved domains in DNA methyltransferase domain-containing proteins were identified using the PFAM database (https://pfam.xfam.org).

### Aspergilli harbor conserved DNA methyltransferases for genome defense and methylation maintenance

The conserved DNA methyltransferase domain is the only domain known to catalyze the addition of methyl groups to DNA specifically at cytosines (5mC) (21). Based on previous domain architecture analyses, phylogeny reconstructions, and functional experiments, five distinct protein families containing a DNA methyltransferase domain have been recognized in fungi: DNMT1, DNMT2, and DNMT5 are commonly found in eukaryotes, while Dim-2 and RID are considered to be fungal specific (21). Among these five families, DNMT2 seems to function as a tRNA methyltransferase rather than DNA methyltransferase (62). In line with previous observation (21,63), DNMT2 only occurs in our study in two Basidiomycota and one Ascomycota (*Schizosaccharomyces pombe*), suggesting it has been lost in most Ascomycota.

Three (DNMT1, DNMT5, and Dim-2) of the four remaining families catalyze the maintenance of DNA methylation pattern from parental DNA strand onto synthesized daughter DNA strands, while Dim-2 is also able to catalyze *de novo* methylation (64,65). Based on our species selection and phylogenetic analyses, we here found *DNMT1* and *Dim-2* to be absent in all Aspergilli and only present in four or seven out of 15 outgroup species, respectively, while *DNMT5* is found in most Aspergilli and outgroup species (Figure 1). Only four outgroup species harboring *DNMT1*, the two Basidiomycota species, *Botrytis cinerea*, and *Plectania melastoma*, which corroborates previous findings (21,66). *Dim-2* was identified only in seven Ascomycota outgroup species, and matches in *B. cinerea* and *Zymoseptoria tritici* were included after an additional TBLASTN search in NCBI (Supplementary Table 6). The original *Dim-2* absence in *B. cinerea* by local Hmmsearch is caused by the older version and fragmented assembly at JGI compared to NCBI. The *Z. tritici* IPO323 strain used in this study carries a non-functional *Dim-2* as previously reported (67) and no significant hit was found with Hmmsearch or BLASTP searches, but several fragmented hits with TBLASTN search in NCBI. This loss of *Dim-2* function in *Z. tritici* IPO323 is very recent because other isolates harbor a functional *Dim-2* gene (67). Its loss leads to increased activity of transposable elements (TEs) and decreased mutation rate, thus influencing genome defense and genome evolution (67). In contrast to the *DNMT1* and *Dim-2* losses in Aspergilli, *DNMT5* is identified in 88 out of 94 Aspergilli and 10 out of 15 outgroup species (Figure 1), with a conserved domain architecture consisting of a DNA methyltransferase domain, an SNF2 family domain (PF00176), and a Helicase conserved C-terminal domain (PF00271), in which the latter two domains are involved in transcription regulation (68). The five outgroup species without DNMT5 are *S. cerevisiae*, *S pombe*, *Neurospora crassa*, *Fusarium graminearum*, and *Cladonia grayi*, among which the two yeast species also lack the other two DNA methylation maintaining enzymes, and thus accordingly are considered to be incapable of DNA methylation (69–71). Five out of the six *Aspergillus* species that lost *DNMT5* belong to a single clade within the *Nidulantes* section (*A. undulatus*, *A. olivicola*, *A. filifera*, *A. venezuelensis*, and *A. stella-maris*), and the sixth one is *A. robustus* which is assigned to the *Circumdati* section (Supplementary Figure 1). In the Basidiomycota fungus *Cryptococcus neoformans*, DNMT5 maintains DNA methylation using the same cofactors as DNMT1 (70,72). Thus, it is most likely that DNMT5 is similarly involved in DNA methylation maintenance in Aspergilli as in *C. neoformans*. Moreover, DNA methylation maintenance does not appear to be required for the survival of fungi as shown with the absence of all three *DNMT1*, *DNMT5*, and *Dim-2* not only within the *Nidulantes* section but also in *S. cerevisiae* and *S. pombe* (Figure 1). However, the absence of these genes could have an impact on the regulation of certain genes like secondary metabolite biosynthetic gene clusters and it would be interesting to compare species within the *Nidulantes* section in more detail to identify how such loss may have affected gene expression and genome organization.

RID is a more recently acquired single domain DNMT that specifically occurs in fungi and is involved in a fungal-specific genome defense mechanism called RIP (Repeat-Induced Point mutation) (21). RID is a putative *de novo* methyltransferase for TEs in both CG and non-CG contexts relating to RIP during the sexual stage in *N. crassa;* however, the capacity of RID to catalyze 5mC has not yet been demonstrated (73). Here, we identified RID in all analyzed Aspergilli species as well as in ten outgroup species. A RID-like homolog knock-out mutant in the model filamentous Ascomycota *Podospora anserina* shows blocked sexual development, as gametes formation and fertilization can happen but dikaryotic cells cannot develop normally (74). Interestingly, the knock-out mutant of *DmtA* (homolog of *RID* in *A. flavus*) shows altered conidiation, sclerotial production, aflatoxin biosynthesis, and virulence (27), suggesting that RID might have additional functions next to genome defense. Additionally, fungi lost the gene for *de novo* DNA methyltransferase, *DNMT3,* around 50–150 million years ago (72,75), and only *DNMT5* and *RID* were consistently found in Aspergilli (Figure 1). Thus, we hypothesize that Aspergilli can only maintain DNA methylation *via* DNMT5 but do not spur genome-wide *de novo* DNA methylation. This hypothesis seems to be corroborated as bisulfite sequencing in *A. flavus* has uncovered very little genome-wide DNA methylation (76). However, systematic analyses of DNA methylation patterns and experimental evidence for the roles of DNMTs in Aspergilli are still largely missing.

### Aspergilli lack *SET7* (*KMT6*), but SET-domain proteins are abundant and other histone methyltransferases are conserved

Histone methylation is typically catalyzed by SET domain-containing proteins (17,77). We identified a total of 1,548 SET domain-containing proteins and classified them into 15 groups based on their domain architecture and phylogenetic support that contain species both from Aspergilli as well as from the outgroup. Among these groups, 107, 111, 109, 102, and 108 sequences represent homologs of the known histone methyltransferases (HMTs) SET9 (KMT5), Dim-5 (KMT1), SET1 (KMT2), Ash1 (KMT2H), and SET2 (KMT3), respectively (Figure 2). The remaining homologs were divided into nine groups that we broadly classified as SET domain-containing proteins. Among them, eight groups contain a SET domain but lack experimentally determined functions, and 117 sequences could be grouped as Ribosomal lysine N-methyltransferases (Figure 2). Enzymes in this group contain a SET domain together with a Rubisco LSMT substrate-binding domain (PF09273). While the homolog in plant transfers a methyl group to the large subunit of the Rubisco holoenzyme complex (78,79), the homolog in *S. cerevisiae* has been characterized as a ribosomal lysine N-methyltransferase (80).

**Figure 2.**
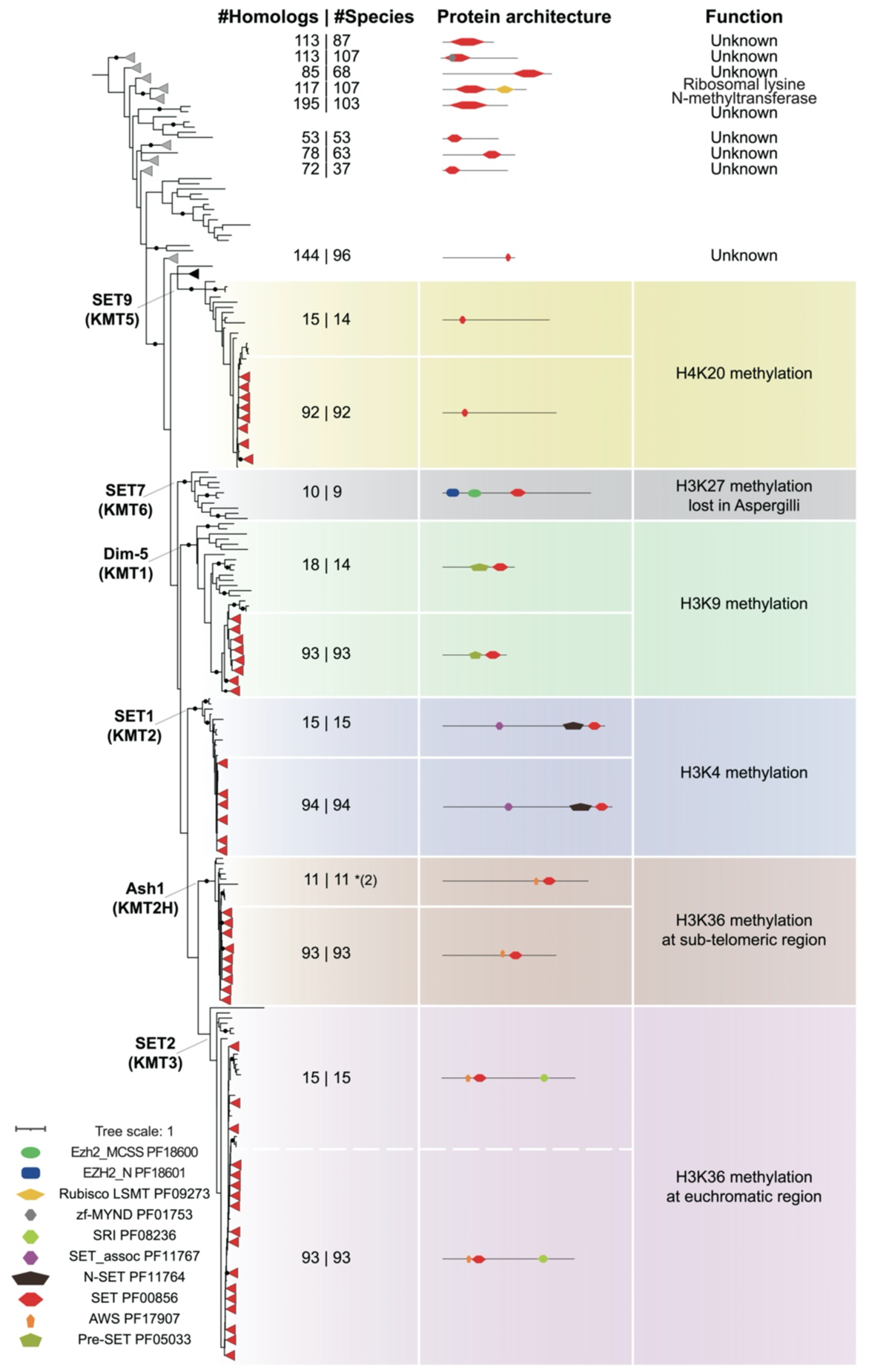
Conservation of histone methyltransferase families in the *Aspergillus* genus and outgroup species. The maximum-likelihood phylogeny of SET domains (PF00856) found in 94 Aspergilli and 15 outgroup species was determined using IQ-TREE (37,38). The black dot on the branch indicates ultrafast bootstrap values over 95 and SH-aLRT values over 80. Leaves from Aspergilli with trustworthy support are collapsed and shown as red triangles. Leaves with similar protein architecture but unknown functions covering both Aspergilli and outgroup species are collapsed and shown as grey triangles. The numbers of hits and species are separated by the pipe in the ‘#Homologs | #Species’ column. The number in parentheses after * indicates the additional hits found by the TBLASTN search on NCBI. Additional conserved domains in SET domain-containing proteins were identified using the PFAM database (https://pfam.xfam.org).

The well-described HMTs, SET1, Dim-5, SET2, Ash1, and SET9, all contain distinct accessory domains and catalyze methylation on lysine residues at different positions of H3 or H4 tails (17). SET1 is composed of the SET domain, the N-SET domain (PF11764), as well as the SET_assoc domain (PF11767), and is responsible for H3K4me1/2/3 in *S. cerevisiae* (81). SET1 occurs in a single copy in all species (Figure 2), indicating that this enzyme and its catalyzed PTM play fundamental roles in fungal biology. Dim-5 contains a SET domain and Pre-SET domain (PF05033) and acts as an H3K9 methyltransferase inducing heterochromatin formation notably in *N. crassa* (82). Dim-5 was found in all species included in this study except *S. cerevisiae* and *A. calidoustus*, among which *S. cerevisiae* is reported to lack H3K9me (82,83). How such a loss has impacted gene expression in *A. calidoustus* in comparison to its closest relative *A. carlsbadensis* deserves further investigation. Both SET2 and Ash1 catalyze H3K36me in *N. crassa* and share the SET domain and the AWS domain (PF17907) (Figure 2), yet SET2 harbor one additional SRI domain (PF08236) that interacts with RNA polymerase II and links H3K36me with transcript elongation (84–86). In *N. crassa,* Ash1 was shown to contribute to H3K36me at sub-telomeric regions, while SET2 acts primarily at euchromatic regions (87,88). Considering the common origin of SET2 and Ash1 as well as their similar activities, an ancient duplication followed by the loss or gain of the SRI domain is likely to have contributed to their functional diversification (Figure 2). *SET2* was found in all outgroup species, while *Ash1* was lost in both Basidiomycota and two yeasts. False absences of *Ash1* in *B. cinerea* and *F. graminearum* were corrected as they could be retrieved by TBLASTN search at NCBI (Supplementary Table 6). Both enzymes are conserved in all *Aspergillus* species except in *A. carbonarius* from the *Nigri* section, suggesting that either other SET domain-containing proteins exhibit functional redundancy or that H3K36 methylation is not fully required in this species. In *Fusarium fujikuroi*, deletion of *SET2* and *Ash1* strongly affected vegetative growth and conidiation (88). SET9 homologs harbor only a SET domain, and were shown to catalyze H4K20me in *S. pombe* (89). This group is well conserved in most of the species except in *S. cerevisiae* (90) and in the two closely related Aspergilli, *A. novoparasiticus* and *A. arachidicola* that both belong to section *Flavi* (Figure 2 and Supplementary Figure 1).

SET7 is the catalytic subunit of PRC2, and is highly conserved in mammals, plants, and *N. crassa*, in which it has been well-characterized to catalyze methylation on H3K27, a PTM that is deposited at facultative heterochromatin (16). Even though PRC2 is considered to be a key chromatin modifier complex, its catalytic subunit *SET7* was lost in many fungal species (17). *SET7* has previously been reported to be absent in *A. nidulans* and *A. fumigatus* (15–17), and we here demonstrate that *SET7* is absent from all analyzed Aspergilli as well as from *Penicillium* (Figure 2), suggesting that this enzyme was most likely lost at or before the divergence of the last common ancestor of these two fungal genera. Thus, in line with previous reports (17,25,91,92), our results indicate that methylation on H3K4, H3K9, H3K36, and H4K20 are likely catalyzed by conserved enzymes in most Aspergilli, while H3K27 methylation is absent. The loss of this methylation is usually reported to trigger serious phenotypes in humans (94) and mice (95). Loss of H3K27me3 in fungi typically leads to gene expression changes without severe growth defects (16,93), and thus further investigating how Aspergilli and other fungi are able to cope with such a loss will likely provide new insights on the histone code and its evolution in fungi.

Importantly, the occurrence of a high number of uncharacterized SET domain-containing proteins opens the possibility that these contribute to not yet characterized histone modifications. Especially, four uncharacterized SET domain-containing groups with 144, 195, 113, and 113 members are conserved in Aspergilli and thus likely play important roles in different cellular processes.

### Histone acetyltransferases are conserved in Aspergilli

Histone acetyltransferases (HATs) have been classified into five families: MYST (MOZ, Ybf2/Sas3, Sas2, and Tip60)-related HATs, Gcn5-related acetyltransferases (GNATs), p300/CBP HATs, general transcription factor HATs, and nuclear hormone-related HATs (96–98). We here only focused on the first two HAT families because these are known to be involved in the regulation of histone proteins and appear largely conserved in fungi (96).The MOZ/SAS FAMILY domain is the catalytic domain of MYST-related HATs (99). As previously reported (100), the three characterized MYST-related HATs, Sas3, Sas2, and Esa1, belong to distinct monophyletic clades, and these paralogues are nearly fully conserved in the here analyzed fungi (Figure 3). MOZ/SAS FAMILY domain and zf-MYST domain (PF17772) are present in Sas3, which can on its own acetylate histone H3 and H4, and H2A weakly *in vitro* (101). Within the NuA3 (Nucleosomal Acetyltransferase of histone H3) complex, Sas3 histone acetylation activity is *in vitro* restricted on H3K14 and to a less extent on H3K23, while in *S. cerevisiae*, only H3K14ac can be detected (102). *Sas3* occurs in most of the species but is lost in *S. pombe*, as previously observed (103,104), and is duplicated in *A. carbonarius* (Figure 3). In addition to the two domains found in Sas3, Sas2 and Esa1 carry a PHD_4 domain (PF16866) or Tudor-knot domain (PF11717), respectively (Figure 3). *Esa*1 is present as a single copy in all species included in this study, and its functions is known to contribute to the acetylation of K5, K8, K12, K16 on histone H4, as well as on H2AZK14 in *S. cerevisiae* (105–108). *Sas2* is similarly found in most of the species except for two Basidiomycota as well as for *A. carbonarius*. Sas2 is involved in chromatin-mediated gene regulation and H4K16 acetylation in both *S. cerevisiae* and *Candida albicans* (105,109,110). Although Esa1 and Sas2 catalyze H4K16ac, it was shown in *C. albicans* that they are differentially recruited at different stages of development (105). Because both Esa1 and Sas2 are well-conserved in the *Aspergillus* genus, we hypothesize that these two HATs may also have distinct roles during development.

**Figure 3.**
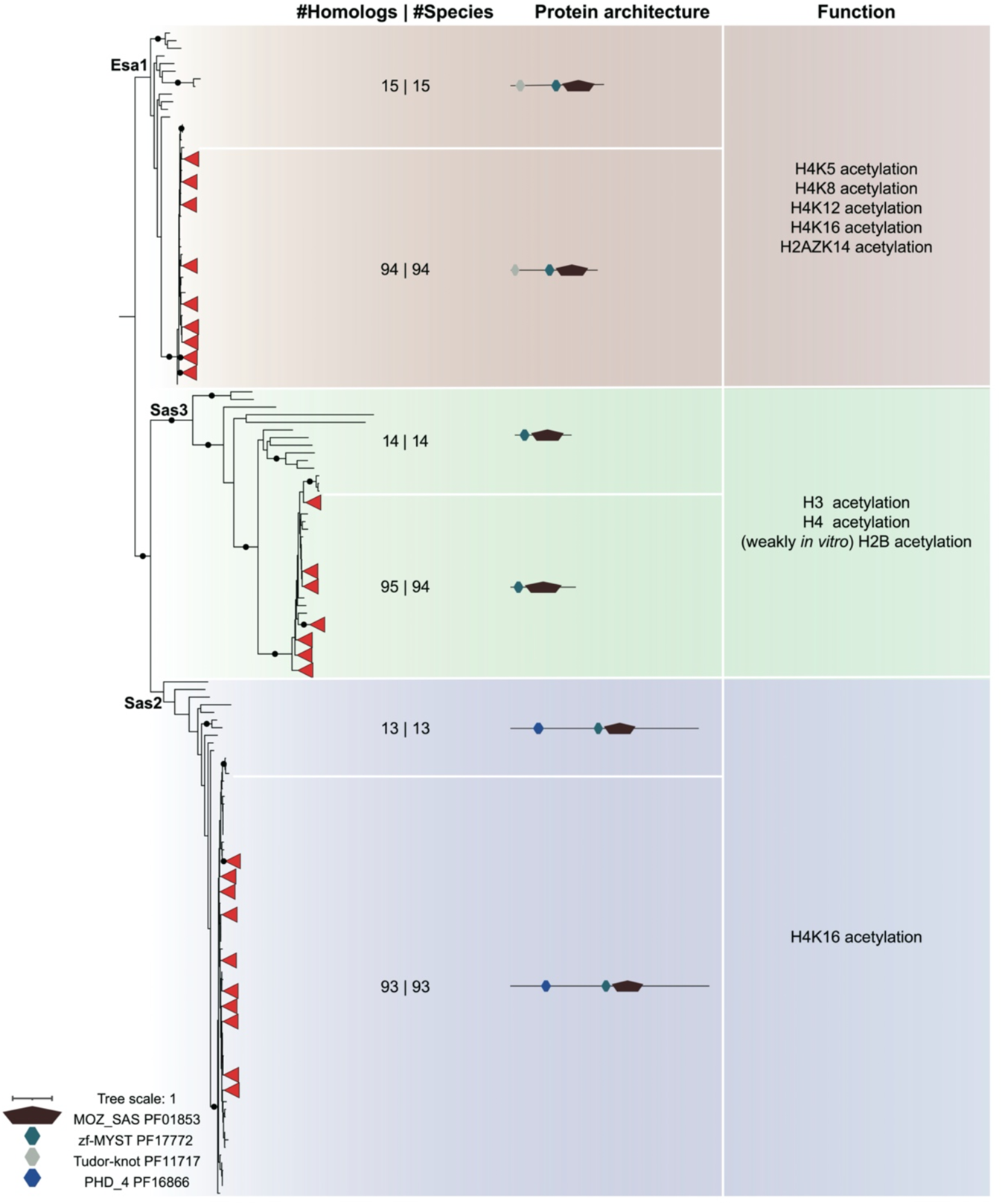
Conservation of MOZ/SAS FAMILY domain-containing histone acetyltransferases in the *Aspergillus* genus and outgroup species. The maximum-likelihood phylogeny of MOZ/SAS FAMILY domains (PF01853) found in 94 Aspergilli and 15 outgroup species was determined using IQ-TREE (37,38). The black dot on the branch indicates ultrafast bootstrap values over 95 and SH-aLRT values over 80. Leaves from Aspergilli with trustworthy support are collapsed and shown as red triangles. The numbers of hits and species are separated by the pipe in the ‘#Homologs | #Species’ column. ‘*In vitro*’ indicates the acetylation activity of this enzyme was only observed on the substrate *in vitro*. Additional conserved domains in MOZ/SAS FAMILY domain-containing proteins were identified using the PFAM database (https://pfam.xfam.org).

GNATs have a acetyltransferase domain as its catalytic core, and 2,604 acetyltransferase domain-containing proteins were found in our search (Supplementary Figure 2). The genes for two well-characterized GNATs, *Gcn5* and *Elp3*, occur in a single copy in every species included in our study (Supplementary Figure 2). Gcn5 harbors a Bromodomain (PF00439) in addition to the Acetyltransferase domain and is a crucial subunit of distinct histone acetyltransferase complexes such as the HAT-A2 complex, the SAGA complex, and the ADA complex in *S. cerevisiae* (96,98). All these complexes contain two coactivator proteins, Ada2 and Ada3, that directly bind to Gcn5 (111) and the recruitment of different other subunits determines the location where they catalyze acetylation (96). Elp3 is characterized by the Acetyltransferase domain, Radical SAM (PF04055), and Radical SAM C-terminal domain (PF16199), and has been shown to catalyze H3K14ac and H4K8ac in *S. cerevisiae* (112–114). Elp3 also exhibits a dual enzymatic activity as it catalyzes demethylation in mouse zygotes (115), which is due to the Radical SAM domain that catalyzes the demethylation of methylated lysyl residues on histones (116).

Next to Gcn5 and Elp3, we have identified 2,386 additional acetyltransferase domain-containing proteins. Because their functions remain unknown due to the lack of homology to sequences with characterized functions, they are valuable candidates that may acetylate histones or other proteins. Interestingly, it has been shown that Gcn5 also exhibits succinyltransferase activity in humans (117) and can also function as histone crotonyltransferase to regulate gene transcription *in vitro* (118). It is expected that other less common histone modifications, i.e., histone lysine butyrylation, propionylation, or malonylation, may also be catalyzed by histone acetyltransferases because these histone modifications all use acyl-Coenzyme A to complete the reactions (119–121). Further research is needed to determine whether these less common histone modifications are catalyzed by the known histone acetyltransferases Gcn5 and Elp3, or as of yet unstudied HATs. In conclusion, all five characterized fungal HATs described above are highly conserved in the *Aspergillus* genus, showing their importance in fungal biology. Many other acetyl-transferase domain-containing proteins are encoded in *Aspergillus* genomes and may be responsible for histone acetylation or additional less common histone modifications like butyrylation, succinylation, and malonylation.

### Duplication of *RpdA* histone deacetylase gene in the *Flavi* and *Janorum* sections

Five families of histone deacetylases (HDACs) can be grouped into two classes based on sequence similarity: class I (HOS1, HOS2, and Rpd3/RpdA) and class II (HdaA and HOS3) (122,123). They all harbor the Histone deacetylase catalytic domain, and our phylogenetic tree corroborates the HDAC classification into two distinct classes (Figure 4). *HdaA* is found as a single copy in each species included in this study (Figure 4). It harbors the Histone deacetylase domain and C-terminal Arb2 domain (PF09757) that serves as an anchor to target centromeric heterochromatin region (124,125). HdaA deacetylates lysine in histones H3 and H2B, but not H4 nor H2A in *S. cerevisiae* (126,127), and the Δ*hdaA* knock-out strains show increased or reduced production of different secondary metabolites in *A. nidulans* (128), *A. niger* (128), and *A. fumigatus* (129). By contrast, *HOS3* is found in only 80 *Aspergillus* species, as it is lost in nine species (*A. nomiae*, *A. pseudocaelatus*, *A. tamarii*, *A. pseudotamarii*, *A. caelatus*, *A. sergii*, *A. novoparasiticus*, *A. minisclerotigenes*, *A. arachidicola*) in the *Flavi* section, three species (*A. desertorum*, *A. stercorarius*, and *A. multicolor*) in the *Nidulantes* section, and two species (*A. heteromorphus* and *A. carbonarius*) in the *Nigri* section. *HOS3* is well conserved in all outgroup species except *S. pombe* (Figure 4) (130). HOS3 has been shown to catalyze deacetylation with distinct specificity for histone H2AK7, H2BK11, H3K14 and H3K23, as well as H4K5 and H4K8 in *S. cerevisiae* cell extracts (131). Notably, *HOS1* was only detected in three outgroup species, *S. cerevisiae* and both Basidiomycota (Figure 4). The specificity of Hos1 in catalyzing deacetylation on histone proteins is not known yet (132), but it can catalyze the deacetylation of the Smc3 subunit of Cohesin and influence the regulation of chromosome segregation during mitosis in *S. cerevisiae* (133). *HOS2* is found in 15 outgroup species and most of Aspergilli except *A. oryzae* (Figure 4). It is known to deacetylate lysine residues in H3 and H4 histone tails (134). The conservation of *HOS2* in Aspergilli suggests that it plays a key role, which is supported by the deletion of *HOS2* in *A. niger* where the deletion strains displays severe reduction in growth, sporulation, SM biosynthesis, and stress resistance (128).

**Figure 4.**
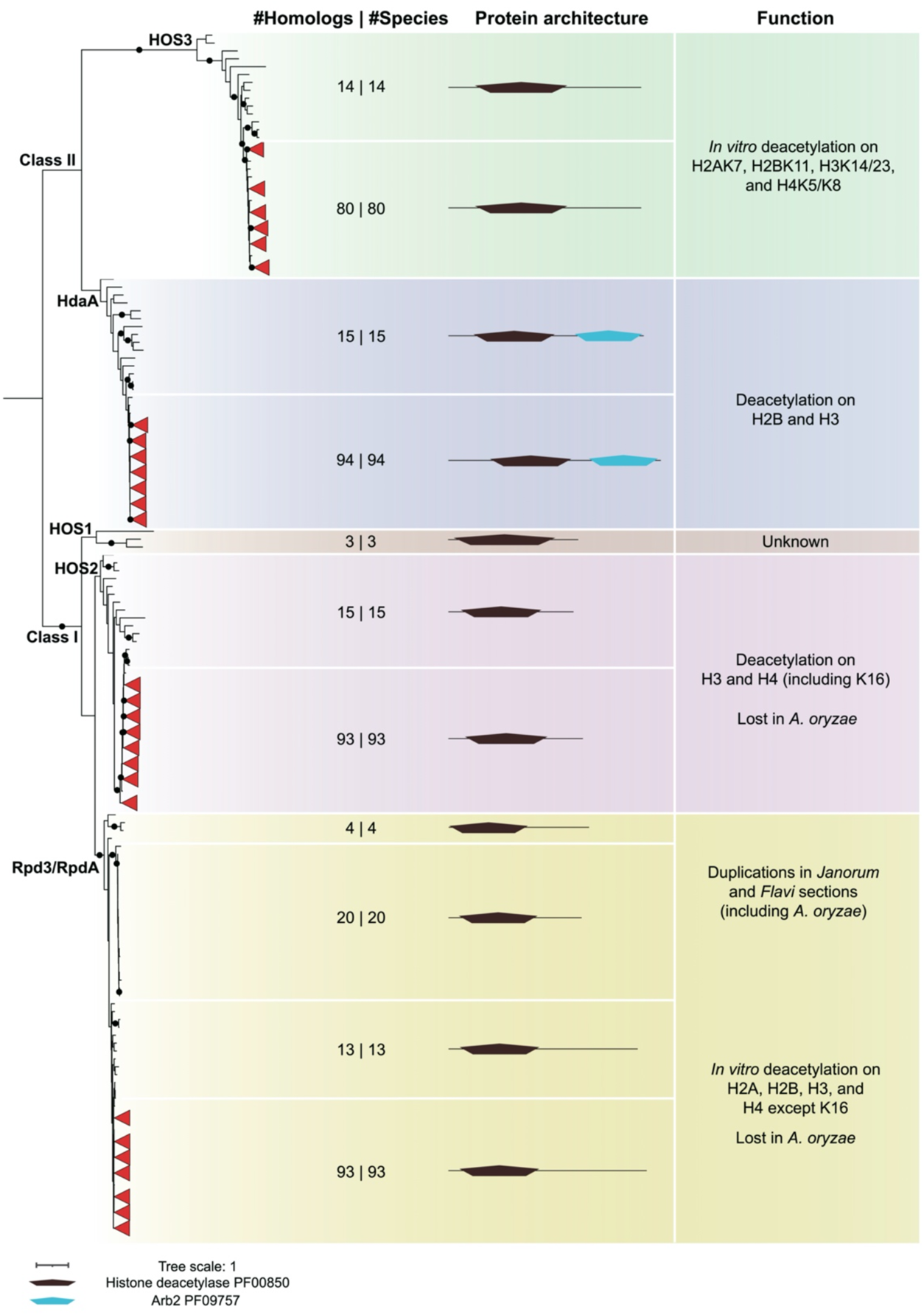
Conservation of histone deacetylase domain-containing proteins in the *Aspergillus* genus and outgroup species. The maximum-likelihood phylogeny of Histone deacetylase domains (PF00850) found in 94 Aspergilli and 15 outgroup species was determined using IQ-TREE (37,38). The black dot on the branch indicates ultrafast bootstrap values over 95 and SH-aLRT values over 80. Leaves from Aspergilli with trustworthy support are collapsed and shown as red triangles. The numbers of hits and species are separated by the pipe in the ‘#Homologs | #Species’ column. ‘*In vitro*’ indicates the deacetylation activity of this enzyme was only observed on the substrate *in vitro*. Additional conserved domains in histone deacetylase domain-containing proteins were identified using the PFAM database (https://pfam.xfam.org).

Within class I HDACs, the Rpd3/RpdA clade is divided into two sub-clades (Figure 4). One is nearly fully conserved (except in *A. oryzae*) and comprises the well-characterized Rpd3/RpdA HDACs, while the other one only comprises sequences from the *Flavi* and *Janorum* sections (Figure 4). Our phylogenetic reconstruction suggests a duplication event that has occurred in the ancestor of all Aspergilli giving rise to a paralog that was retained in two sections only and could explain the *A. oryzae* loss in the conserved sub-clade. The two paralogs from the same species vary in length as they share a conserved N-terminal catalytic domain but have a different C-terminus (Figure 4). Follow-up studies are needed to investigate the function of both paralogues in *Aspergillus* species. Rpd3 is a crucial deacetylase in eukaryotes, as it is required for deacetylation in most locations on core histone proteins, except H4K16 *in vitro* (135) that is catalyzed by HOS2 (134). *S. cerevisiae* lacking *Rpd3* is sensitive to high osmolarity and shows compromised expression of osmotic stress genes (136). Deletion of *RpdA* in *A. nidulans* is lethal (137), while only heterokaryon disruptants could be obtained in *A. oryzae* (138), indicating its crucial role in the fungal genus *Aspergillus*. In summary, class I and class II deacetylases are well conserved in Aspergilli, and duplication events like the one observed for *RpdA* may confer new activities. While only one paralog was retained in *A. oryzae*, the presence of two paralogs in sections *Flavi* and *Janorum* may provide functional redundancy or diversification, which remains to be determined.

Lastly, we reconstructed the phylogeny of histone demethylases harboring the Jumonji C (JmjC) domain (PF02373) (Supplementary Figure 3). Six groups are distinguished by their domain distribution: KDM4A/JHDM3/JMJD2, KDM5/JARID, DMM-1, JMJD6/PKDM11, KDM2/Epe1, and KDM6B/JMJD3, which also agrees with a previous study (91). The first five groups occur in most of the outgroup and *Aspergillus* species, but the *KDM6B/JMJD3* is only present in several outgroup species but not in any of the *Aspergillus* species. Certain JmjC domain-containing proteins were shown to catalyze the demethylation of histone proteins, e.g., KDM5/JARID for H3K4me2/3 and KDM4A/JHDM3/JMJD2 for H3K9me2/3 and H3K36me2/3 (139). The histone demethylase activity has not been detected for DMM-1 (140) or Epe1 (141) in *N. crassa*, while KDM2 can demethylate H3K36me1/2 in mouse (142) and JMJD6/PKDM11 is an arginine demethylase in *Arabidopsis* (143). Interestingly, the human KDM6B/JMJD3 is responsible for the demethylation of H3K27me2/3 (144). As we did not find *SET7* and thus conclude that H3K27me3 is absent, it would be tempting to speculate that H3K27 demethylase should be absent in *Aspergillus* too. However, based on the fact that other enzymes, e.g., UTX can also catalyze the demethylation of H3K27 (144,145), we cannot unambiguously correlate the absence of *KDMB6B/JMJD3* to *SET7* loss. More studies are needed to investigate the regulation mechanism and interplay of proteins for H3K27 methylation.

### Presence-absence patterns of subunits that compose histone modifier complexes

Histone PTMs are catalyzed by the conserved domains found in HMTs, HATs, and HDACs (Figure 2–4, Supplementary Figure 2), but these chromatin modifying enzymes are often not acting alone. Instead, they are part of protein complexes that comprise accessory subunits, which are involved in the recognition and induction of specific histone modifications at the correct locations (13,14). To further investigate the presence-absence patterns (PAPs) of known chromatin modifier complexes in Aspergilli, we focused on 15 chromatin modifier complexes with 83 subunits previously identified and studied in *S. cerevisiae* or *N. crassa* (Supplementary Table 4). Forty-four subunits are present in all *Aspergillus* and outgroup species. Most of the accessory subunits are either fully conserved or absent in Aspergilli (Figure 5 and Supplementary Table 4). Ten subunits show PAPs in both Aspergilli and outgroup species, while nine subunits appear specific to *S. cerevisiae* as they are not found in any other species and are likely to play a specific role in this species. Seventeen subunits show PAPs in the outgroup species, while eight are absent and nine are present in the Aspergilli. Lastly, three subunits are present in all outgroup species but show PAPs in Aspergilli.

**Figure 5.**
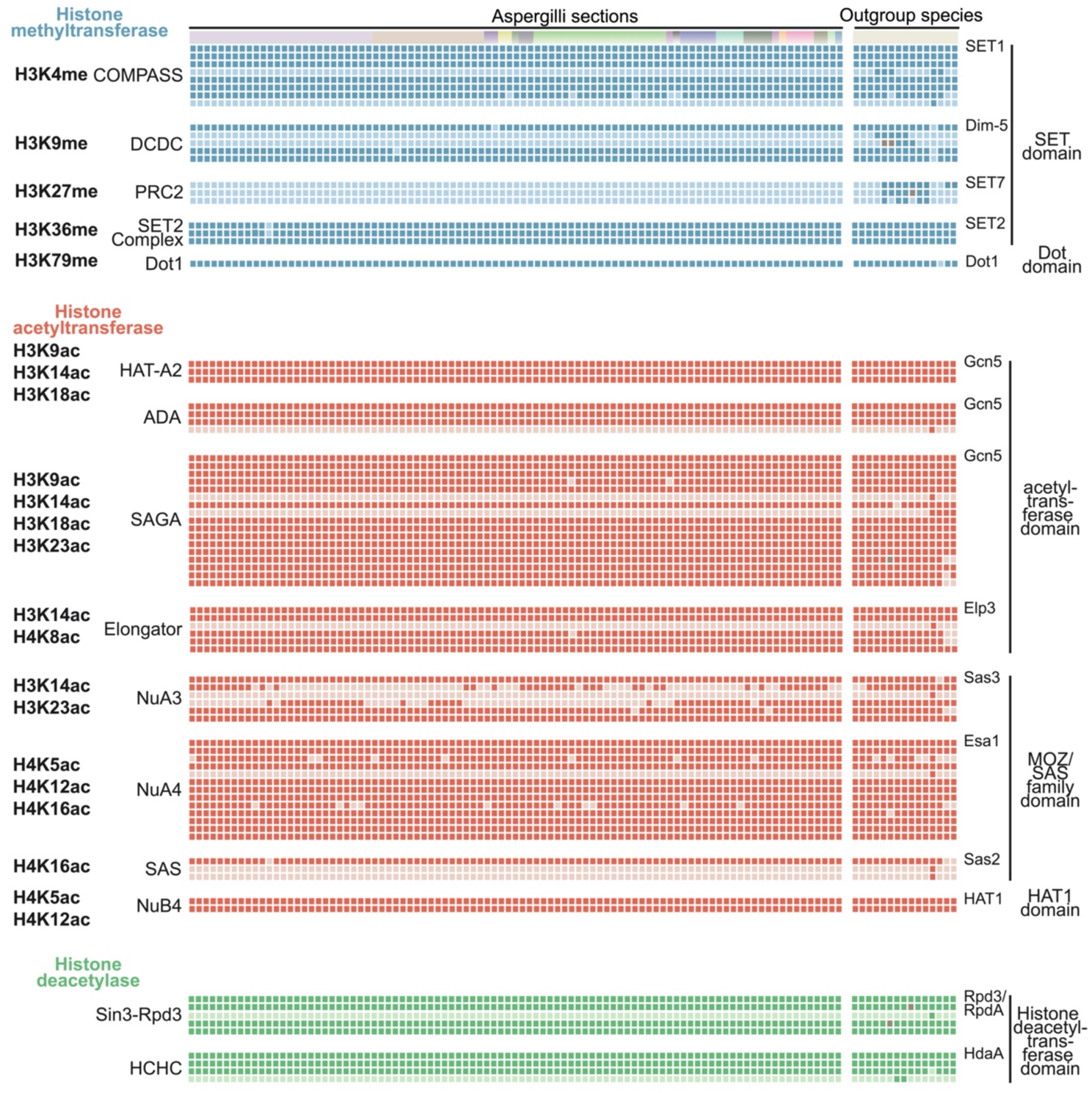
Presence-absence patterns of subunits that constitute characterized histone modification complexes. The top row with distinct color boxes indicates outgroup species and Aspergilli sections, the order of the species from right to left is the same as in the species tree from top to bottom (Supplementary Figure 1) without *Fusarium oxysporum* and *Aspergillus kawachii* since they were excluded in the later analysis. Each row represents a subunit in the histone modification complexes, the order agrees with the summary list (Supplementary Table 4). Each row shows the presence/absence of a specific complex subunit. For each complex, the first row represents the catalytic subunit. Each box in the matrix indicates the presence (dark color) or absence (light color) of a subunit (Supplementary Table 4). The brown boxes indicate additional hits found with TBLASTN search on NCBI (Supplementary Table 5).

Five histone methyltransferase complexes, COMPASS, DCDC (Dim-5/-7/-9/CUL4/DDB1 Complex), SET2 Complex, PRC2, and Dot1 (disruptor of telomeric silencing 1) (KMT4) were included in our analyses (Figure 5). Five COMPASS subunits (*SET1*, *CclA*, *Swd1*, *Swd2*, and *Swd3*) are fully conserved, suggesting an important function for H3K4 methylation. Consistent with a PAP in most outgroup species and the loss in Aspergilli, Spp1p is not essential for H3K4 methylation (146). As *Sdc1* shows a PAP in Aspergilli and outgroup species, it is likely that this subunit also does not play a key role in H3K4 methylation. In the DCDC complex, while Dim-7 and Dim-9 seem to be dispensable for H3K9 methylation, Dim-5, Cul4, and Dim-8 are highly conserved, and we thus expect these enzymes to play key roles in the activity of this complex (Figure 5 and Supplementary Table 4). However, it was shown that Dim-7 and Dim-9 are responsible for heterochromatin recognition, which leads to subsequent recruitment of the complete DCDC to induce H3K9 methylation in *N. crassa* (147). Thus, it is very likely that other accessory subunits are involved in recruiting DCDC at specific chromatin locations in Aspergilli. Furthermore, the unique absence of *Dim-8* in *A. indicus* may suggest altered H3K9 methylation in this species. Consistent with the absence of *SET7* (Figure 2), all components of the PRC2 complex are lost in Aspergilli (Figure 5). Lastly, the interaction and crosstalk of multiple chromatin modifications in *Aspergillus* species should be further explored. For example, heterochromatin formation and regulation are done by H3K9me3, DNA methylation, and H3K27me3 in *N. crassa*. In this species, H3K9me3 is recognized by an adapter protein, HP1, to recruit DNA methyltransferase Dim-2 for DNA methylation (73), and cause the redistribution of H3K27me3 (15). In *Aspergillus* species, although HP1 and the writer for H3K9me3, DCDC complex, are well conserved (Figure 5), the inability for genome-wide *de novo* DNA methylation and H3K27me3 suggest different mechanisms of heterochromatin formation and regulation in Aspergilli which remain to be determined.

We included eight histone acetyltransferase complexes that differ by their distinct catalytic subunits (Figure 5). The HAT-A2, ADA, SAGA, and Elongator complexes use an acetyltransferase domain-containing protein as catalytic subunit (29,148). The SAS (Something About Silencing), NuA4 (Nucleosomal Acetyltransferase of histone H4), and NuA3 complexes use a MOZ/SAS FAMILY domain-containing protein (149). The NuB4 (Nuclear Hat1p-containing type B histone acetyltransferase) complex uses the histone acetyltransferase HAT1 domain-containing protein (150). Gcn5, Ada2, and Ada3 are fully conserved in all fungi and constitute the core enzymatic module of the HAT-A2, ADA, and SAGA complexes. This catalytic module contributes to catalyzing histone acetylation, and other subunits in these complexes guide them to specific locations and link them to the transcriptional machinery (151). Besides the catalytic module, the ADA complex also harbors the Ahc1 subunit that is found in *S. cerevisiae* only (148,152). Twelve additional subunits of the SAGA complex are well conserved in Aspergilli and outgroup species (Figure 5). The SAGA complex also contains a deubiquitylase (DUB) module, for which the composition differs between Aspergilli and other species. *Sgf11* is unique to *S. cerevisiae* and *Sus1* was likely lost in the ancestor of Pezizomycotina because we can only identify *Sus1* orthologs in both Basidiomycota and the yeasts (Figure 5). The Elongator complex appears more variable as four subunits are mostly conserved, but *Elp6* is specific to *S. cerevisiae*, and *Hpa2* and *Hpa3* show PAPs in both Aspergilli and outgroup species.

The NuA4, SAS, and NuB4 complexes are well-conserved in Aspergilli as well as in the outgroup species (Figure 5). These complexes appear slightly different in *S. cerevisiae* with the presence of specific subunits (*Eaf5* in the NuA4 complex; *Sas4* and *Sas5* in the SAS complex). Both SAS and NuA4 complexes can catalyze H4K16ac with opposite effects during cell growth and morphogenesis in *C. albicans* (105). It would be interesting to assess whether these two complexes exhibit similar activities in Aspergilli. For the NuA3 complex, two accessory subunits are well conserved among Aspergilli while two others, *Pdp3*, and *Nto1*, show different PAPs, and *Yng1* is specific to *S. cerevisiae*. *Pdp3* and *Nto1* seem to be lost in a complementary manner, suggesting a possible functional redundancy (Figure 5). Finally, two histone deacetylase complexes, Sin3-Rpd3 and HCHC, are fully conserved (Figure 5). The Sin3-Rpd3 complex is slightly different in *S. cerevisiae* with the specific of the *Ume1* subunit.

Overall, our analysis showed that the core catalytic modules of complexes involved in histone PTMs are mostly conserved in all fungi, including Aspergilli, but variability in accessory subunits could contribute to differences in the localization of histone PTMs along the genome.

### Unbiased mass spectrometry captures the diverse histone modifications in *A. nidulans* and verifies the absence of H3K27 methylation

Our genomic and phylogenetic analyses have identified the presence of multiple chromatin modifiers in Aspergilli, suggesting that these species have the ability to establish a wide variety of histone modifications. To assess the presence of histone PTMs predicted in our evolutionary analyses (Fig 2–5), we extracted histone proteins from *A. nidulans* and performed unbiased mass spectrometry analyses (Figure 6A and 6B). H3 and H4 peptides with PTMs corresponding to H3K4me, H3K9me, H3K36me, and H3K79me were detected (Figure 6C), which we anticipated being catalyzed by SET1, Dim-5, Ash1 and SET2, and Dot1, respectively. H4K20me, which is catalyzed by SET9 in *S. pombe* (89), was not detected (Figure 6C), which is likely due to technical reasons because the unmodified form of histone H4 peptide 20-24 containing the K20 residue was not detected. For the acetylation on H3K9, H3K14, H3K18, H3K23, H4K5, H4K8, H4K12, and H4K16, they were confidently detected (Figure 6C). We did not observe any peptide that would indicate the presence of H3K27me in *A nidulans*, while we did detect H3K27ac (Figure 6C and Supplementary Table 6). These results corroborate our computational analysis that the PRC2 complex is absent in *A. nidulans*, as well as previous reports on the absence of H3K27me3 in several Aspergilli (17,153,154). Thus, the mass spectrometry results verify the occurrence of histone modifications predicted to be present based on our comparative analyses in the fungal genus*Aspergillus*. Moreover, the differences between the relative abundance of distinct histone modifications provide interesting information to investigate these abundances and identify specific conditions under which these modifications increase or decrease in abundance.

**Figure 6.**
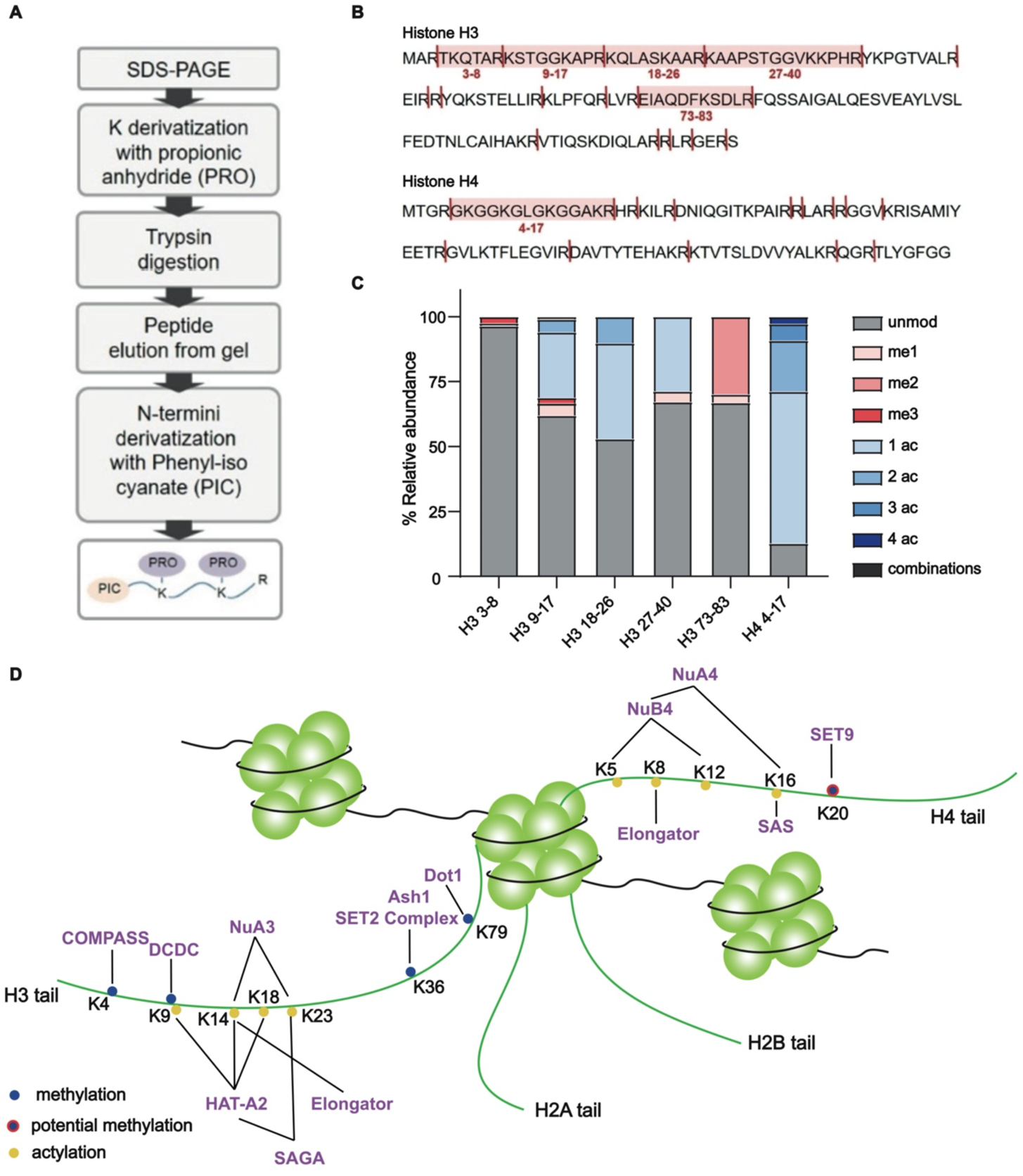
Mass spectrometry unlocks the histone code in Aspergilli. **A.** Schematic presentation of the in-gel digestion method used to process histones isolated in *A. nidulans*. After SDS-PAGE separation, gel bands corresponding to histone H3 and H4 are excised and chemically acylated with propionic anhydride (which derivatizes unmodified and mono-methylated lysines). Thanks to the derivatization, trypsin cutting at lysines is impaired, and histones are cleaved only at the C-terminus of arginine. Finally, digested peptides are N-terminally derivatized with phenyl isocyanate (PIC). **B.** *A. nidulans* histone H3 and H4 sequences. Cleavage sites after PRO-PIC digestion are shown. Peptides identified in this study are highlighted in light red and labeled using their starting-ending amino acids. **C.** Relative abundance of post-translational modifications detected on peptides of histone H3 and H4 from *A. nidulans* using unbiased mass spectrometry analysis. unmod: unmodified peptide; me1: monomethylation; me2: dimethylation; me3: trimethylation; ac: acetylation. The term “combinations” refers to multiple modifications occurring on the same peptide (Supplementary Table 6). **D.** Histone modifications are predicted by genomic and phylogenetic analyses, and/or proved by mass spectrometry in *A. nidulans*. Histone modifications verified by mass spectrometry are labeled as solid dots, while the ones that are only predicted by phylogenic analysis are labeled as solid dots with a red outline.

## Conclusion

Changes in chromatin organization play an important role in regulating the expression of genes (130). The fungal genus *Aspergillus* is well known for its ability to thrive in a broad range of different environmental niches and the huge potential of secondary metabolite biosynthesis (11,155). However, a detailed and complete overview of occurrence and evolution of chromatin modifiers in Aspergilli is a prerequisite to be able to better understand how these fungi regulate gene expression. In the future, more research is needed to uncover how the absence of *de novo* DNA methylation impacts genome evolution in Aspergilli. We demonstrate that PRC2 is lost throughout Aspergilli, a finding consistent with the lack of H3K27me detection by mass spectrometry. Although histone acetyltransferases and deacetylases are overall conserved, our study also revealed specific duplication and loss events, which require further investigation to understand their impact on chromatin and gene regulation in certain *Aspergillus* species. Moreover, other less common histone modifications like histone lysine succinylation, crotonylation, butyrylation, propionylation, and malonylation, are all likely catalyzed by HATs and HDACs as they all use Coenzyme A (CoA) to complete the reactions (119–121), but may require specific yet to be discovered subunits. In conclusion, our study provides a detailed overview of the evolutionary routes of chromatin modifier complex in the fungal genus *Aspergillus*, and therefore generates the necessary framework to identify targets for functional studies to understand how chromatin regulates gene expression in Aspergilli.

## Acknowledgment

The sequence data were produced by the US Department of Energy Joint Genome Institute (http://www.jgi.doe.gov/) in collaboration with the user community. Especially, we thank the Scott Baker, Mikael Andersen, and Joseph Spatafora for their permission to use the sequence data. We thank Jos Houbraken for validating the *Aspergillus* species tree and Joseph Strauss for providing *A. nidulans* strain. We thank Alessandro Vai for support with analysis of mass spectrometry data for histone PTMs’ identification on *Aspergillus* core histone sequences.

## Authors and contribution

X.Z. formal analysis, investigation, methodology, visualization, writing original draft; R.N. formal analysis, investigation, methodology, visualization; T.B. funding acquisition, supervision, resources; J.C. conceptualization, funding acquisition, methodology, project administrations, supervision, visualization, writing original draft; M.F.S. conceptualization, funding acquisition, methodology, project administrations, supervision, visualization, writing original draft.

## Supplementary materials

**Supplementary Figure 1. A robust species phylogeny for the fungal genus *Aspergillus*.** The species phylogeny was based on a maximum-likelihood phylogeny reconstructed by IQ-TREE with partitioned analysis using 758 BUSCO (Benchmarking Universal Single-Copy Orthologs) genes. The dot on the branch indicates ultrafast bootstrap values over 95 and SH-aLRT bootstraps values over 80. Species from the same section are labeled by background color with a section name. Two species, *Fusarium oxysporum* and *Aspergillus kawachii*, are included in the species tree, but we don’t use these two in the following analyses.

**Supplementary Figure 2. Two known histone acyltransferase-domain containing protein groups are conserved in the Aspergillus genus and outgroup species together with many unknown proteins.** Maximum likelihood phylogeny of Histone acyltransferase domains (PF00583) found in 94 Aspergilli and 15 outgroup species were determined using IQ-TREE (Chernomor et al. 2016; Nguyen et al. 2015). The black dot on the branch indicates ultrafast bootstrap values over 95 and SH-aLRT bootstraps values over 80. Additional conserved domains in histone acyltransferase domain-containing proteins were identified using the PFAM database (https://pfam.xfam.org).

**Supplementary Figure 3. Five histone demethylase-domain containing protein groups are conserved in *Aspergillus* genus and outgroup species while KDM6B/JMJD3 is lost.** Maximum likelihood phylogeny of Jumonji C (JmjC) domain (PF02373) found in 94 Aspergilli and 15 outgroup species were determined using IQ-TREE (Chernomor et al. 2016; Nguyen et al. 2015). The black dot on the branch indicates ultrafast bootstrap values over 95 and SH-aLRT bootstraps values over 80. Additional conserved domains in JmjC domain-containing proteins were identified using the PFAM database (https://pfam.xfam.org).

**Supplementary Table 1. List of species included in this study.**

**Supplementary Table 2. List of BUSCO genes used to build the species tree.**

**Supplementary Table 3. List of sequence names in the domain trees.**

**Supplementary Table 4. List of histone modifier complexes reported in the literature.**

**Supplementary Table 5. Identification of missing chromatin modifiers in the outgroup species using TBLASTN search**

**Supplementary Table 6. Mass spectrometry result for *A. nidulans*.**

**Supplementary Data 1. Fasta, trimmed alignment, and maximum-likelihood phylogenetic tree files associated with Figure 1–4 and Supplementary Figure 2-3.**

**Supplementary Data 2. Fasta, trimmed alignment, and maximum-likelihood phylogenetic tree files associated with Figure 5.**

