## Supplementary material for "The histone code of the fungal genus *Aspergillus* uncovered by evolutionary and proteomic analyses": Figure S1

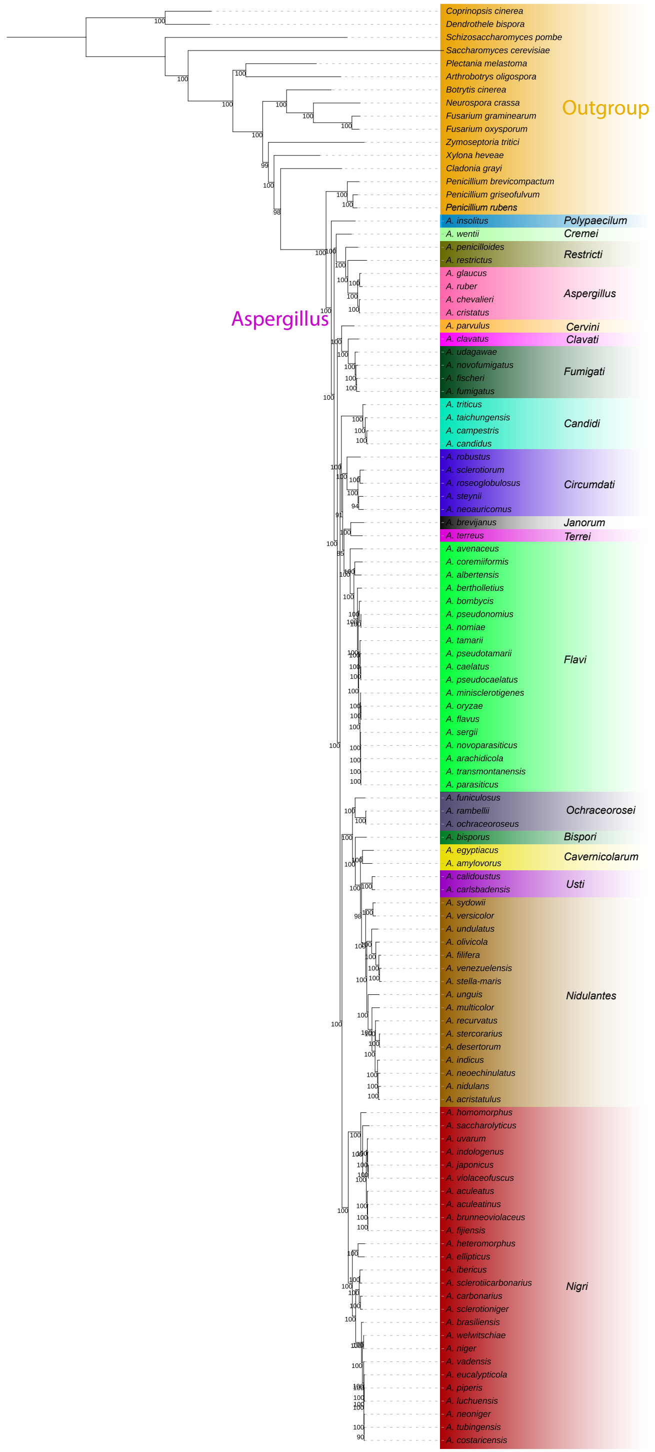


**Supplementary Figure 1. A robust species phylogeny for the fungal genus *Aspergillus*.** The species phylogeny was based on a maximum-likelihood phylogeny reconstructed by IQ-TREE with partitioned analysis using 758 BUSCO (Benchmarking Universal Single-Copy Orthologs) genes. The dot on the branch indicates ultrafast bootstrap values over 95 and SH-aLRT bootstraps values over 80. Species from the same section are labeled by background color with a section name. Two species, *Fusarium oxysporum* and *Aspergillus kawachii*, are included in the species tree, but we don't use these two in the following analyses.
