## Supplementary material for "The histone code of the fungal genus *Aspergillus* uncovered by evolutionary and proteomic analyses": Figure S2

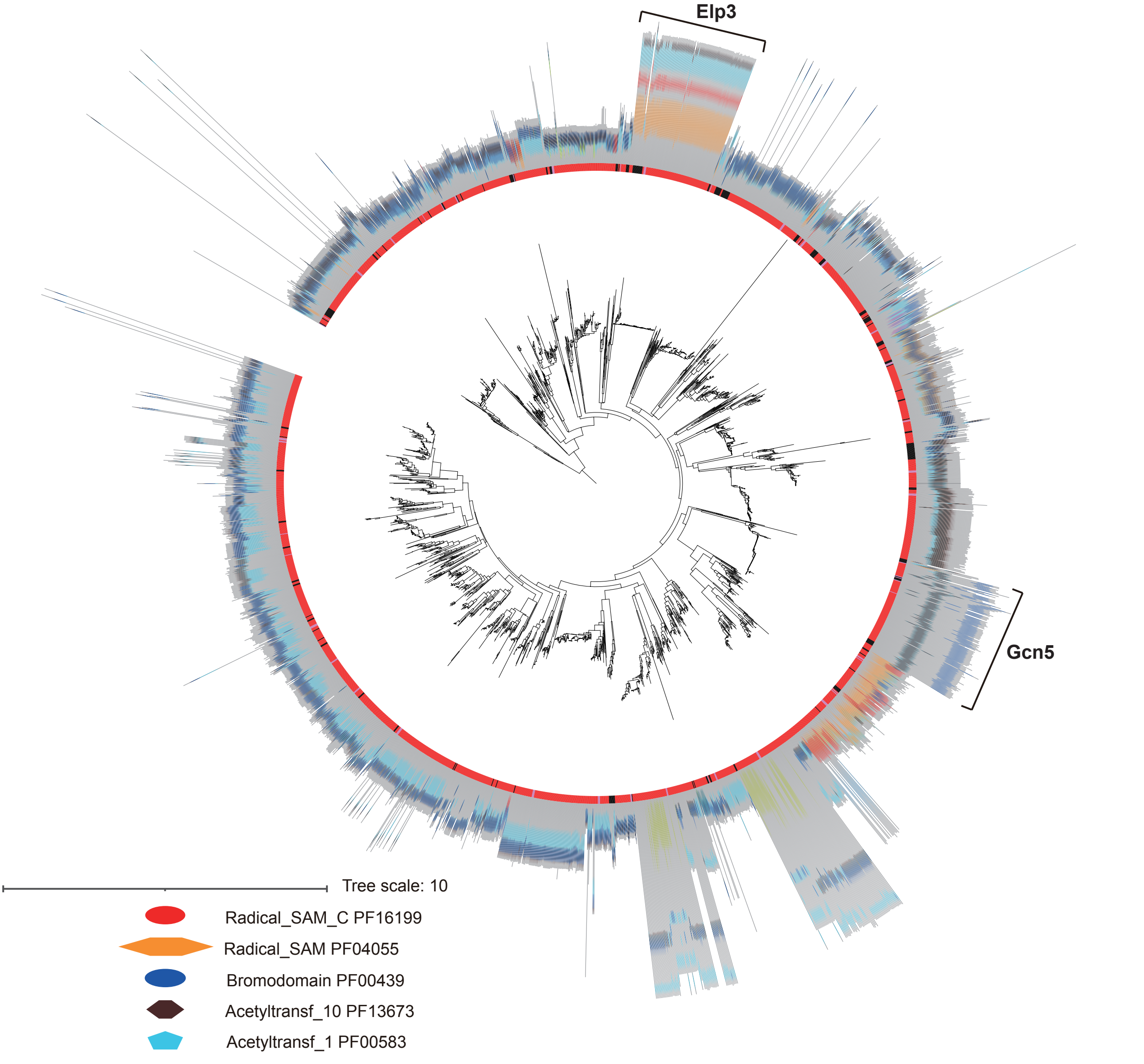


**Supplementary Figure 2. Two known histone acyltransferase-domain containing protein groups are conserved in the *Aspergillus* genus and outgroup species together with many unknown proteins.** Maximum likelihood phylogeny of Histone acyltransferase domains (PF00583) found in 94 Aspergilli and 15 outgroup species were determined using IQ-TREE (Chernomor et al. 2016; Nguyen et al. 2015). The black dot on the branch indicates ultrafast bootstrap values over 95 and SH-aLRT bootstraps values over 80. Additional conserved domains in histone acyltransferase domain-containing proteins were identified using the PFAM database (https://pfam.xfam.org).
