## Supplementary material for "The histone code of the fungal genus *Aspergillus* uncovered by evolutionary and proteomic analyses": Figure S3

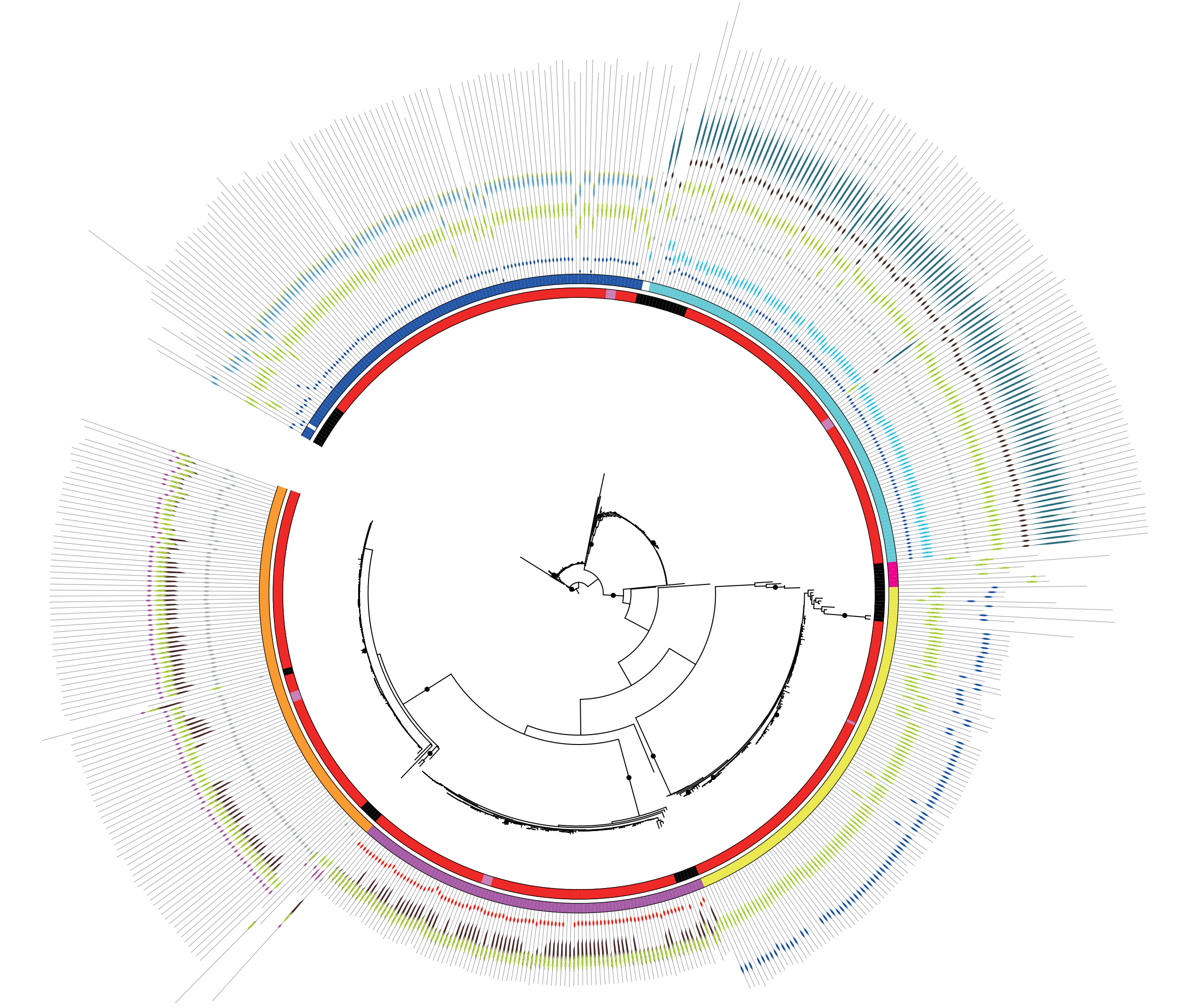

Tree scale: 1

- JmjC PF02373
- zf-HC5HC2H PF13771
- JHD PF17811
- Cupin_8 PF13621
- PLU-1 PF08429
- zf-C5HC2 PF02928
- zf-HC5HC2H_2 PF13832
- ARID PF01388
- PHD PF00628
- JmjN PF02375
- F-box-like PF12937
- zf-4CXXC_R1 PF10497

| Group | Function |
|---|---|
| **KDM4A/JHDM3/JMJD2** | Demethylate H3K9/K36me2/3 |
| **Unknown** | Unknown |
| **KDM5/JARID** | Demethylate H3K4me2/3 |
| **KDM6B/JMJD3** | Demethylate H3K27me2/3 in human |
| **DMM-1** | Demethylase activity is not detected yet in *Neurospora crassa* |
| **JMJD6/PKDM11** | Demethylate H4R3me2 in *Arabidopsis* |
| **KDM2/Epe1** | Demethylate H3K36me1/2 in mouse, no demethylase activity detected in *Neurospora crassa* |

**Supplementary Figure 3. Five histone demethylase-domain containing protein groups are conserved in *Aspergillus* genus and outgroup species while KDM6B/JMJD3 is lost.** Maximum likelihood phylogeny of Jumonji C (JmjC) domain (PF02373) found in 94 Aspergilli and 15 outgroup species were determined using IQ-TREE (Chernomor et al. 2016; Nguyen et al. 2015). The black dot on the branch indicates ultrafast bootstrap values over 95 and SH-aLRT bootstraps values over 80. Additional conserved domains in JmjC domain-containing proteins were identified using the PFAM database (https://pfam.xfam.org).
